# Simultaneous CRISPR screening and spatial transcriptomics reveals intracellular, intercellular, and functional transcriptional circuits

**DOI:** 10.1101/2023.11.30.569494

**Authors:** Loϊc Binan, Serwah Danquah, Vera Valakh, Brooke Simonton, Jon Bezney, Ralda Nehme, Brian Cleary, Samouil L Farhi

**Author notes:** Corresponding authors: Brian Cleary, Samouil L Farhi.

## Abstract

Pooled optical screens have enabled the study of cellular interactions, morphology, or dynamics at massive scale, but have not yet leveraged the power of highly-plexed single-cell resolved transcriptomic readouts to inform molecular pathways. Here, we present Perturb-FISH, which bridges these approaches by combining imaging spatial transcriptomics with parallel optical detection of *in situ* amplified guide RNAs. We show that Perturb-FISH recovers intracellular effects that are consistent with Perturb-seq results in a screen of lipopolysaccharide response in cultured monocytes, and uncover new intercellular and density-dependent regulation of the innate immune response. We further pair Perturb-FISH with a functional readout in a screen of autism spectrum disorder risk genes, showing common calcium activity phenotypes in induced pluripotent stem cell derived astrocytes and their associated genetic interactions and dysregulated molecular pathways. Perturb-FISH is thus a generally applicable method for studying the genetic and molecular associations of spatial and functional biology at single-cell resolution.

## Introduction

Pooled loss of function screening has proven to be a powerful application of CRISPR-Cas9 technology, identifying the genetic underpinning of phenotypes such as enhanced growth^1^, drug resistances^2^, and virus infection rates^3^. The scalable experimental workflow of pooled guide RNA (gRNA) cloning, lentiviral packaging, and subsequent sequencing-based gRNA readout allows for high throughput experiments where large numbers of genes are perturbed. More recently, pooled screening has been combined with single cell RNA-sequencing (scRNA-seq) to combine suppression of target genes with transcriptome wide expression measurements in the same cell^4–6^. This has allowed mapping gene circuits within cells and determination of causal connections between the expression of a gene and that of downstream pathways. However, these approaches share the limitations of dissociated scRNA-seq measurements: namely, that the cell’s local context is lost. Thus, though it is known that gene expression in a given cell is affected by its direct neighborhood^7,8^, these effects cannot be determined from droplet based CRISPR screens. To decipher intercellular genetic interactions, one needs to be able to record gene expression and gRNA perturbations in their spatial context.

Optical pooled screening provides an alternative to sequencing based pooled screens by combining *in situ* gRNA sequencing with imaging-based profiling. These methods are extremely high throughput but have not been linked to gene expression measurements^9–11^. Incorporating spatial transcriptomic measurements of *in situ* gene expression profiles would allow not only probing gene regulatory circuits at both intra- and inter-cellular level, but also pairing such data with functional measurements of the system, such as time-lapse recording of the response of cells to a given stimulus. Dhainaut et al. have demonstrated a combination of gRNA perturbation and sequencing-based spatial transcriptomics in Perturb-map^12^, by using protein barcodes of gRNAs paired with 10x Visium to spatially readout perturbation effects. While this approach successfully pairs transcriptomics, CRISPR perturbation, and spatial information, the use of protein barcodes sacrifices the advantages of pooled cloning workflows, and the spatial resolution of Visium limits the use of the approach on cells that do not grow clonally. There is thus still a need for a tool that combines the benefits of single-cell spatial measurements with *in situ* gRNA read out at scale.

To meet this need, we turned to imaging spatial transcriptomics (iST) for its resolution, efficiency, scalability, and compatibility with home-brew reagents at low cost. However, any technology that probes both transcriptome and gRNAs using microscopy needs to address a major challenge: identifying single molecules over background autofluorescence, which requires signal amplification. Multiplexed Error Robust Fluorescence In Situ Hybridization (MERFISH) achieves this for mRNA measurements by tiling each mRNA with a minimum of 25 “encoding” probes, each of them annealing on a 30 base pairs (bp) region of the target molecule^13,14^. These barcoded encoding probes are then read out over multiple rounds of staining, imaging, and destaining to recover mRNA identity. The barcodes are structured with a Hamming weight of 4, meaning that individual molecules light up in 4 of the images acquired, and Hamming distance of 4, meaning that errors in binding can be identified and corrected computationally. Although this strategy yields single molecule detection, gRNAs cannot be tiled in the same way as they are only 20bp long. MERFISH was previously used in conjunction with CRISPR perturbations using a long barcode and a reporting gene^15^, but this approach is limited by the challenges of pooled cloning of long barcodes; lentivirus packaging efficiency of long insert sequences; and lentiviral shuffling of barcodes distant from the gRNA^10,16^ In addition, the branched amplification readout strategy used in this work was noisy and required the expression of a reporter RNA, further complicating the experimental implementation. We reasoned that to leverage barcoded MERFISH readouts while allowing pooled workflows on gRNAs, gRNAs can be used as self-barcodes if adequately amplified *in situ*. Rather than tiling, we used an alternative approach for amplification by generating many local copies of the target molecule, by *in situ* transcription via T7 polymerase^17^ (**Fig 1**).

**Figure 1.**
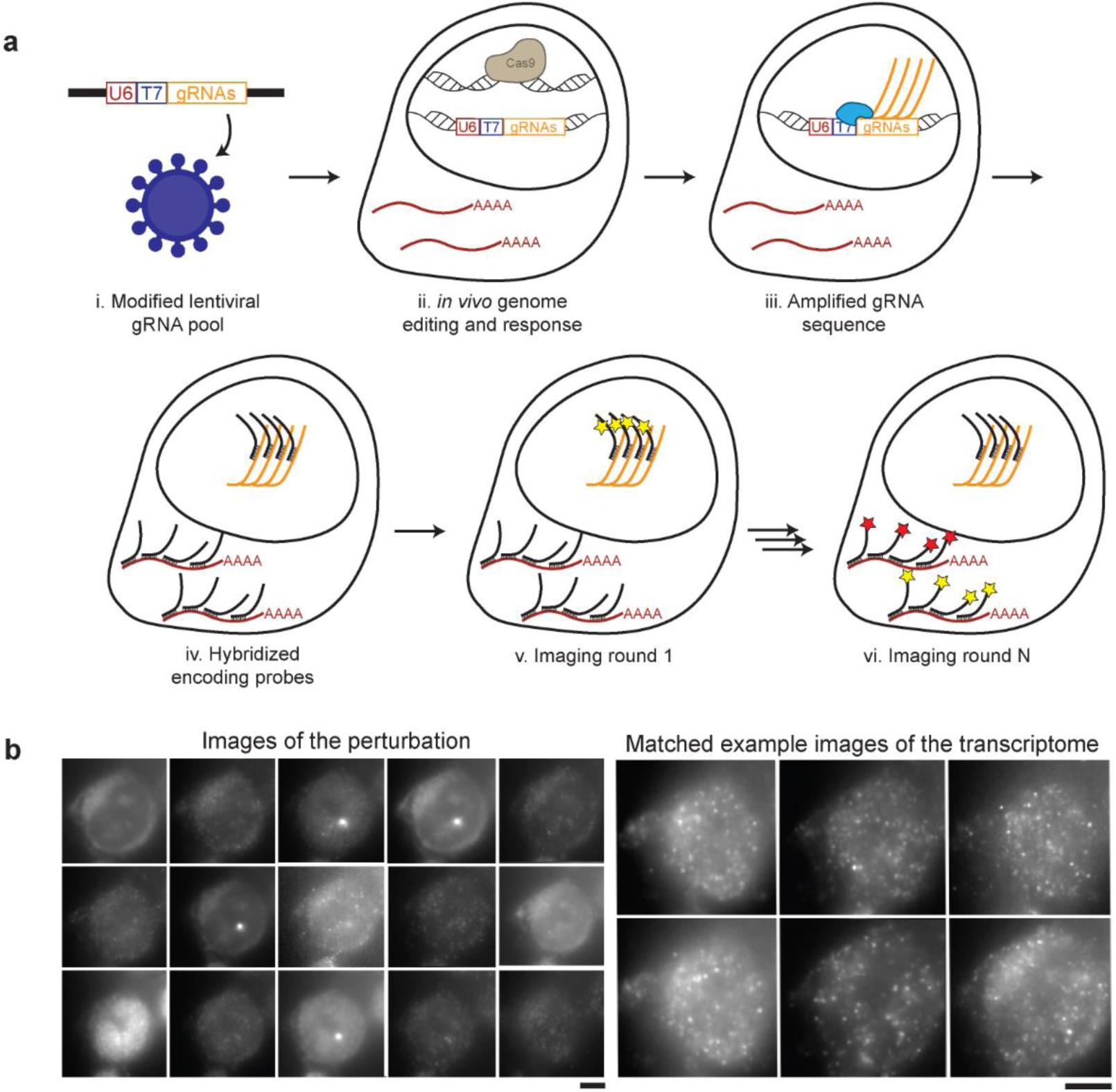
Perturb-FISH allows recording both gRNAs and transcriptome in cells in their spatial context. A) Perturb-FISH workflow. i) Guides are packaged into lentiviral particles using a modified version of Lentiguide-puro that contains a T7 promoter between the end of the U6 promoter and the beginning of the guide. ii) The lentivirus is used to insert the guide sequence in the genome of the cell, resulting in genome editing. iii) T7 polymerase locally generates many copies of the gRNA in fixed cells. iv) DNA encoding probes anneal on both the target mRNA and the amplified gRNA. v) Fluorescent readouts anneal on the encoding probes, and are imaged. The fluorophores are cleaved, and this step is repeated. vi) After sequentially imaging the gRNAs, rounds of hybridization/imaging/cleavage continue to image the transcriptome in the same cells. B) Representative images of Perturb-FISH: Amplified gRNA generate a bright spot in the nuclei of cells, and the identity of gRNAs is encoded in the sequence of images in which they fluoresce (left). The transcriptome is read out the same way with MERFISH (right). T7 transcription yields higher signal amplification than the tiling of an mRNA with 30 probes, as visible by the larger size of the spots they generate. Scale bar = 10μm.

Here, we introduce Perturb-FISH, which combines MERFISH with local amplification of the gRNA region to decode both perturbations and transcriptome in their spatial context. We demonstrate Perturb-FISH in a system for which we had previously generated matched data using Perturb-seq (**Fig 2a**): the genetic network regulating the response of THP1-derived macrophages to lipopolysaccharide (LPS) stimulation^18^. We validated Perturb-FISH by comparing the CRISPR knock out (KO) effects of gRNA perturbation as recovered by Perturb-FISH with those recovered by Perturb-seq (**Fig 2b**). We then demonstrated the power of Perturb-FISH to interrogate intercellular genetic circuits by examining how cell density (**Fig 2c**) impacts the effects of gRNA perturbations and how cells are affected by the presence of certain perturbations in their direct neighbors. Finally, we extended Perturb-FISH by coupling a functional readout in a CRISPR inhibition (CRISPRi) screen of autism spectrum disorder (ASD) risk genes. We record knock down (KD) effects on both calcium activity (assayed via live imaging) and gene expression in human induced pluripotent stem cell (hIPSC) derived astrocytes (**Fig 2d**).

**Figure 2.**
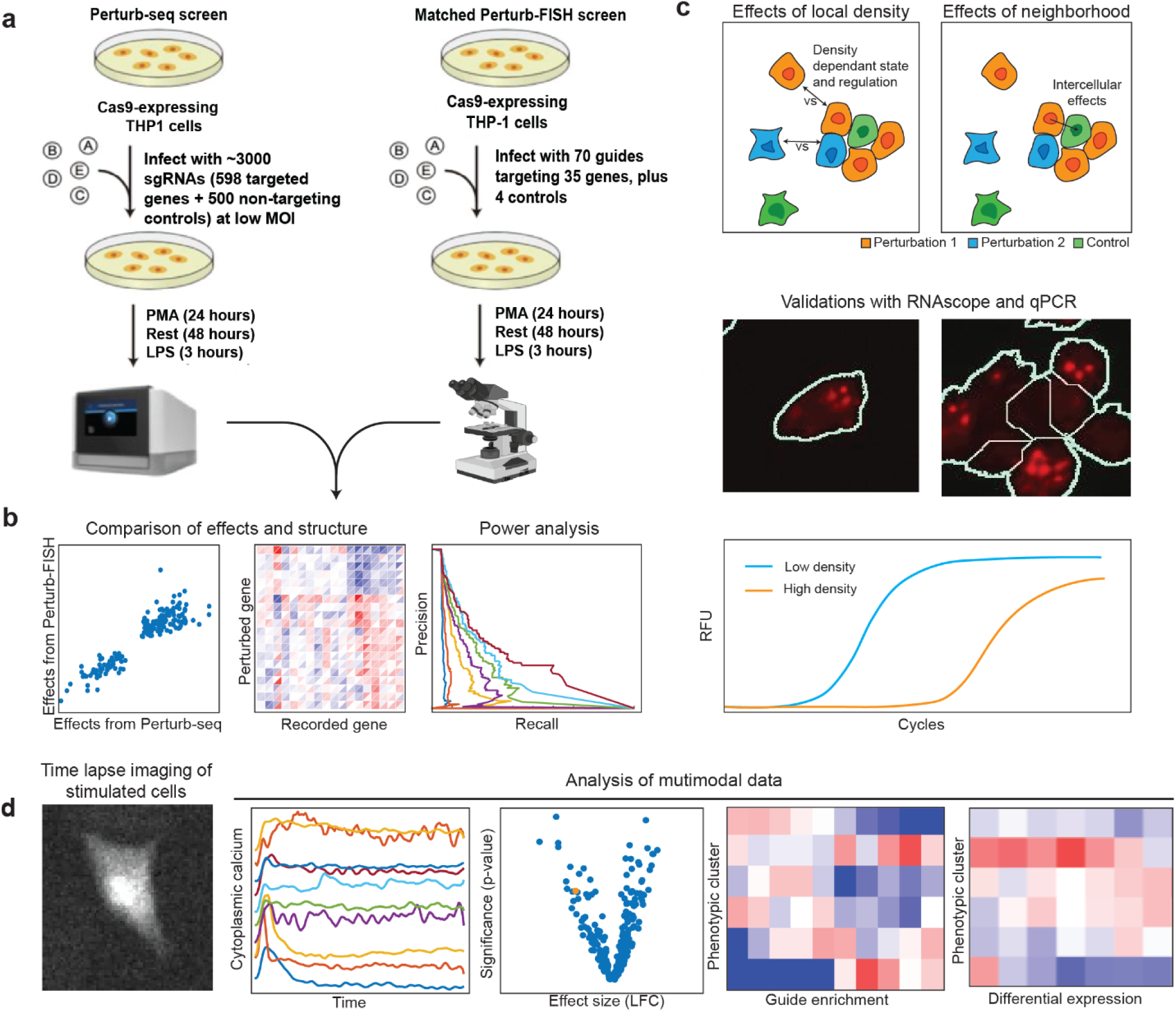
Overview of experimental design and analysis. A) Overview of matched Perturb-seq and Perturb-FISH screens for genetic regulators of LPS response in THP1-cells. B) We compare perturbation effect sizes from the two screens to evaluate Perturb-FISH in terms of consistency, recapitulation of global regulatory structure, and power (Fig 3). C) We first characterize the effects of local cell density on perturbations effects, then we recover effects from perturbed cells onto their unperturbed control neighbors. We validate these results using both qPCR and RNAscope (Fig 4). D) We demonstrate the use of Perturb-FISH in conjunction with live imaging characterization of calcium phenotypes, in a screen of ASD risk genes (Fig 5).

## Results

### Perturb-FISH allows recording both gRNA and mRNA *in situ*

We modified a lentiviral vector commonly used to transduce cells with gRNAs (lentiguide-puro)^19^ to include a T7 phage promoter between the end of the U6 promoter (responsible for the expression of gRNAs in the live cell) and the beginning of the gRNA sequence itself (**Fig1 a.i**). As shown by Romanienko et al.^20^, this fused U6T7 promoter retains its capacity to express functional gRNAs *in vivo* through activation of the U6 promoter, while also allowing *in situ* amplification in fixed cells using the T7 promoter. Because transcription from the U6 promoter starts 23 bases after the end of the essential sequence, the T7 sequence is not transcribed with the guide. Local amplification of DNA from a T7 promoter is used in the ZOMBIE (Zombie is Optical Measurement of Barcodes by In situ Expression) protocol^17^ to allow decoding single cell barcodes *in situ*.

In Perturb-FISH, this *in situ* transcription protocol (**Fig 1a.iii**) is followed by a proteinase based clearing step and a standard MERFISH protocol involving staining with a pre-designed encoding probe library containing probes against both guides and mRNAs of interest (**Fig 1a.iv**). During imaging, 20 bp fluorescent imaging, or “recording,” probes are sequentially flowed onto the sample and imaged^13,14^ (**Fig 1a.v** and **Fig 1a.vi**). Between each round, the fluorophore is cleaved off from the readout probes by disulfide reduction. To make all chemistries compatible, incubation temperatures for both gRNA and mRNA encoding probes must be similar. We found that the 20 bp gRNA length was too short: the melting temperature of 20 bp gRNA encoding probes was lower than the temperature of the formamide washes that are part of the MERFISH protocol. We extended the target region into the first ten bases of the gRNA scaffold, which allowed us to record multiplexed data for both gRNAs and mRNAs. High multiplexing is achieved using the same approach as MERFISH to encode the identity of each perturbation: each RNA molecule is assigned a sequence of 4 (in this case out of 15) images in which it is “on” (**Fig 1b**). Computational reconstruction registers images, assigns barcodes to locations, performs error correction built into the library design, and finds gRNA and mRNA identities versus spatial locations (**Methods**). The images are then segmented to yield count tables of mRNA expression and gRNA identity.

### Perturb-FISH recovers genetic perturbation effects with high correlation both intrinsically and with Perturb-seq

We first applied Perturb-FISH to a system for which we have matched data from Perturb-seq: the response of THP1-derived macrophages to (LPS) stimulation^18^. Cells were perturbed using a library of 74 gRNAs against 35 target genes, differentiated into macrophages, and then stimulated with LPS (**Fig 2a**) After images were decoded and the data condensed in count tables, the effect of each perturbation was computed using FR-perturb, a method we recently developed to infer perturbation effects^18^. We generated replicate datasets from two coverslips to assess reproducibility.

The inferred effects were self-consistent within Perturb-FISH experiments and highly concordant with Perturb-Seq results. Effects of perturbations for key genes (*MAP3K7, IRAK1, TRAF6, RELA*) of the NFκB pathway were highly correlated between the two Perturb-FISH samples (Pearson correlation coefficient of significant effects from Perturb-FISH replicates of 87, 93, 90, 85% respectively), with all statistically significant effects across perturbations being 92% correlated (**Fig 3a** and **Supplementary figure 1a**). Significant effects for those four genes are also highly correlated between each Perturb-FISH replicates and the matched Perturb-seq data, with respective Pearson correlations of 87, 78, 82 and 75% between sample 1 and Perturb-seq, and 68, 79, 87 and 80 between sample 2 and Perturb-seq (**Fig 3b**). Since data from both Perturb-FISH were highly correlated, we pooled them together to increase statistical power for the rest of the study. After pooling, the correlation between pooled Perturb-FISH data and Perturb-seq data remains high at 79% for effects significant in both modalities. As expected from the overall agreement in individual perturbation effects, the global structure of perturbation effects is largely consistent between modalities: both extract the same modules of target genes that affect cells in a similar way when perturbed, as well as comparable programs of genes that react similarly to a given perturbation (**Fig 3c**). These clusters match previous knowledge of the system with *TNF*, *TNFAIP2* and *IL1a* being among the genes most downregulated by perturbing targets such as *MAP3K7*, *IRAK1*, *TRAF6*, *RELA*, *MYD88*^18,21,22^.

**Figure 3.**
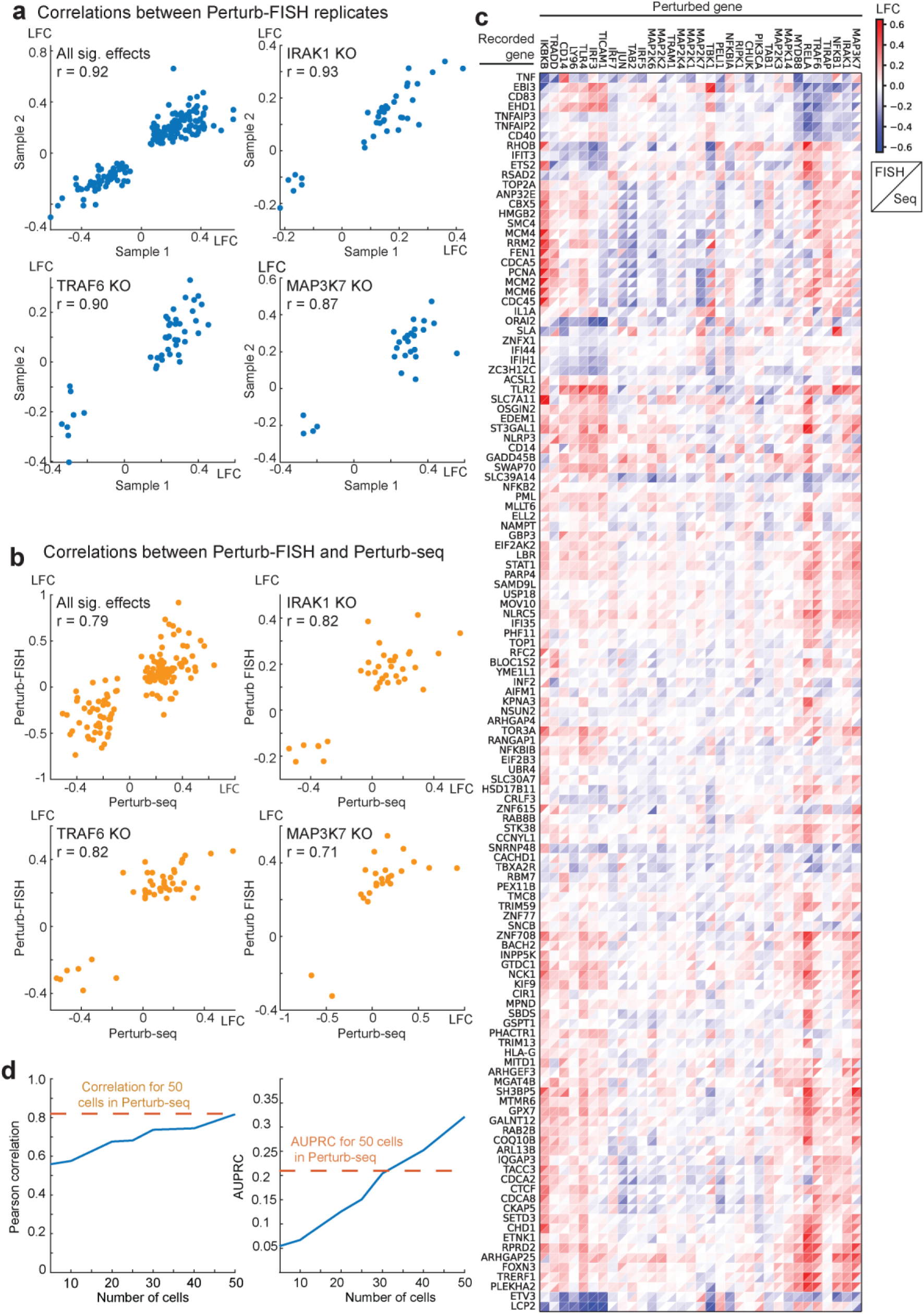
Perturb-FISH robustness and power analysis. A) Scatterplots of significant effect sizes (q<0.1) determined in Perturb-FISH sample 1 (x-axis) and the same effect sizes as determined in sample 2 (y-axis). Effects represent log-fold changes (LFC; natural log base) in expression relative to control cells. Individual plots depict significant effects across all perturbations or for individual perturbations MAP3K7, IRAK1 and TRAF6. B) Scatterplots of significant effect sizes (q<0.1) determined in Perturb-seq (x-axis) and combined analysis of both Perturb-FISH replicates (y-axis). Effects represent log-fold changes (LFC; natural log base) in expression relative to control cells. Individual plots depict significant effects across all perturbations or for individual perturbations MAP3K7, IRAK1 and TRAF6. C) Heatmap of LFC effect sizes (colorbar) found in Perturb-FISH (upper left triangle of each square) and the same effects in Perturb-Seq results (lower right triangle). Rows and columns are clustered based on UPGMA clustering of Perturb-FISH data. D) Held out validation accuracy (y-axis; Pearson correlation, left; AUPRC, right) of effects learned in downsampled Perturb-FISH data using increasing numbers of cells (x-axis). Orange line shows approximate correlation (left) or AUPRC (right) obtained with 50 cells in Perturb-seq, estimated using previously published data^18^.

While testing the consistency of Perturb-FISH data, we noticed a few outlier effects that correlate between Perturb-FISH datasets but do not correlate with Perturb-seq (**Supplementary Fig 1b**). These come from perturbations *IRF3*, *TLR4*, and *CD14* that showed no effect in Perturb-seq, but gave significant effects in Perturb-FISH. Notably, significant effects from these perturbations are highly concordant between Perturb-FISH datasets, and they agree with the current understanding of the pathway as the knock-out of the main LPS receptor *TLR4* is expected to downregulate genes from the immune response like *TNF* or *IL1a*^23,24^. Similarly, *CD14* KO^25^ and *IRF3* KO^26^ were both shown to upregulate *TNF* expression in response to LPS.

Next, we ran a power analysis to identify the minimal number of cells per target to get reliable effects by Perturb-FISH. To do so, we reserved half of the data as a validation set, and used the other half to test the consistency of the inferred effects as we increased the number of cells used for each perturbation, from 5 to 50 cells. Plotting only the effects that are statistically significant according to the validation set, we observed that correlation of effects sizes between subsampled test set and validation set steadily increases with the number of cells used in the smaller subset, from 68% with 20 cells to 74% with 30 and 40 cells, and finally reaching 81.7% with 50 cells, which is comparable to what we achieved with our matched Perturb-Seq study (**Fig 3d**). For all sizes, considering only the effects that are statistically significant according to both the test set and the validation set gives correlations above 98%. Finally, we built precision/recall curves and calculated area under the precision/recall curve (AUPRC) (**Fig 3d**, and **Supplementary Fig 1c**) showing improved precision and recall (starting around 30 cells) compared with matched Perturb-seq data^18^. Therefore, Perturb-FISH allows making hypotheses on perturbations effects with power comparable to Perturb-Seq using 20-50 cells per perturbation target.

### Perturb-FISH recovers cell extrinsic effects of gene perturbation

The previous data highlight that Perturb-FISH recovers the intracellular effects of genetic perturbations at least as well as Perturb-seq. We next investigated the additional information that is afforded by an *in situ* approach, in particular the effects of local cell density on gene expression. When grown on a glass surface, THP1s will spontaneously organize in areas of variable cell density (**Fig 2c** and **Supplementary Figure 1d**). We observed that the expression of certain genes has a high dependency on cell density. In particular, expression of hallmark LPS response genes *TNF* and *IL1a* varied considerably with the number of immediate neighbors to a cell (**Fig 4a**), with *TNF* raw counts increasing up to 1.34-fold with the number of neighbors and *IL1a* raw counts decreasing up to 2.6-fold. The expression levels of *TNF* and *IL1b* have been previously reported to depend on cell density, although the directionality of the effect is inconsistent between studies^27–30^.

**Figure 4.**
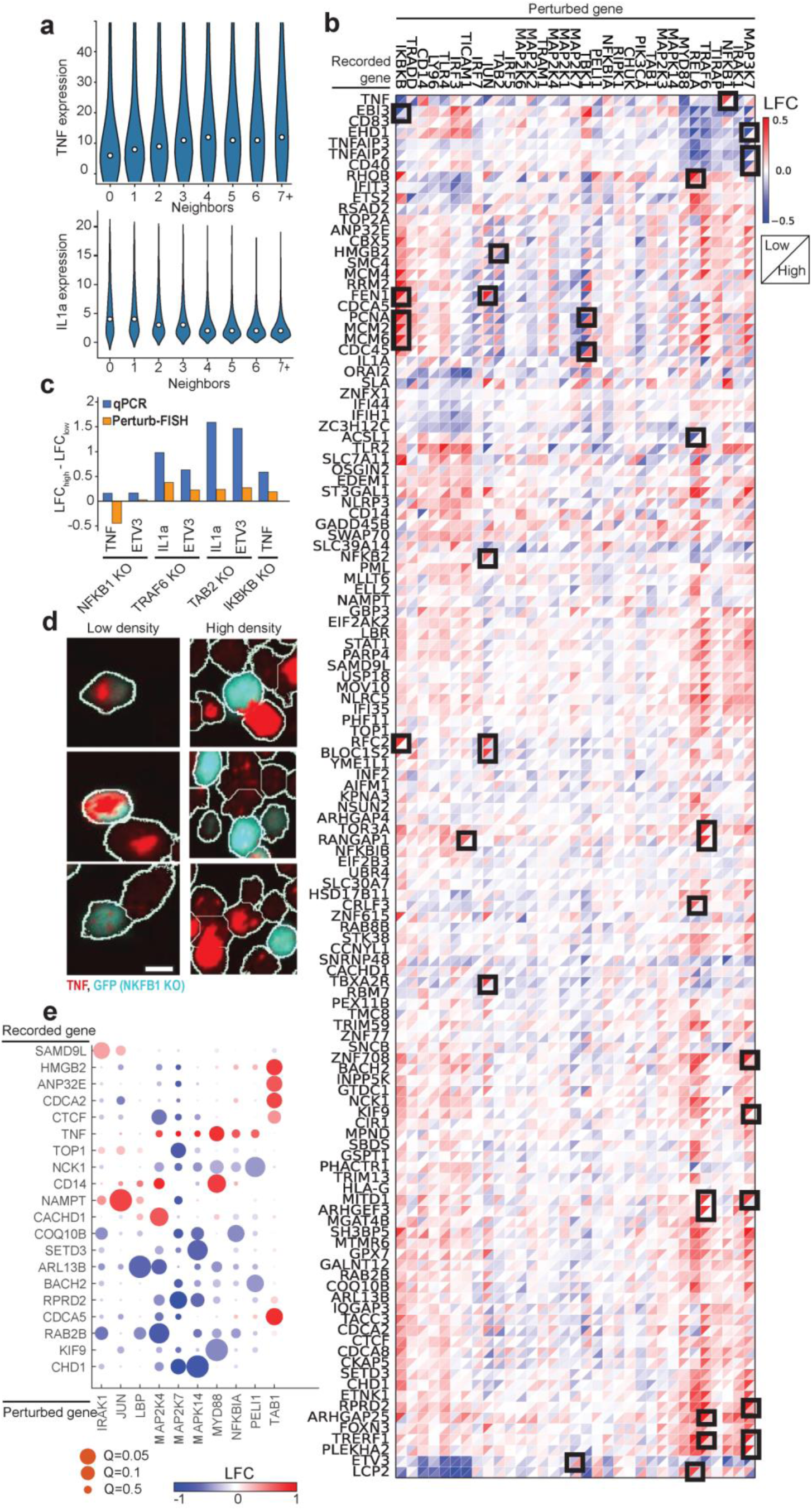
Perturb-FISH analysis of density-dependent and intercellular effects. A) Violin plots show the expression level (y-axis) of TNF (top) and IL1a (bottom) in cells with different numbers of neighbors (x-axis). B) Heatmap of LFC effect sizes (colorbar) found by Perturb-FISH in cells with 2 or fewer neighbors (upper left triangle of each square) or 3 or more neighbors (lower right triangle). Effects represent log-fold changes (LFC; natural log base) in expression relative to control cells at matching density. Black squares highlight effects that are significant (q<0.1) in at least one density, and different by at least 0.4 between densities. C) Comparison of the density dependency of the effects of perturbing NFkB1, TRAF6, TAB2 or IKBKB on TNF, ETV3 or Il1a expression. Bars show the difference between effects at high density and effects at low density, when effects are measured with Perturb-FISH (in orange), or with a qPCR (blue). Effects are expressed as log-fold changes (LFC; natural log base). D) RNAscope images showing TNF transcripts (red) in NFkB1 KO cells (blue) and unperturbed cells (black) at both low density (left) and high density (right). E) Heatmap of LFC effect sizes (colorbar) of gene knock outs (x-axis) on the expression of genes (y-axis) in neighboring unperturbed cells. Effects are expressed as log-fold changes (LFC; natural log base). Circle size shows significance, with q values from 0.05 to 0.5.s

In light of these results, we analyzed the Perturb-FISH data to identify density-dependent effects of genetic perturbation. To do so, we split the data into one set of cells at low local density (with two or fewer contacting neighbors) and another set of cells at high density (three or more contacting neighbors). While dividing the datasets brought the analysis close to the limit of the statistical power of this dataset, we discovered that while most perturbations showed concordant effects at the two densities, many perturbations have substantially different effects depending on cell density (**Fig 4b**). Of note, *NFKB1* KO results in an upregulation of *TNF* only in cells at low density (log fold change at low density (LFC_low_): 0.47, Q-value: 0.014, LFC_high_: 0.03). Meanwhile, *TRAF6* KO results in a much stronger *IL1a* downregulation at low density (LFC: −0.44, Q-value: 0.14) than it does at high density (LFC; −0.06, Q-value: 0.7) (**Fig 4b**). *ETV3* is another example of gene that is further downregulated at low density compared with high density by both *TRAF6* KO (LFC_low_: −0.38, LFC_high_: −0.17) and *NFKB1* KO (LFC_low_: −0.53, LFC_high_: −0.27). Note that LFCs in this text use a natural log basis.

We used two approaches to validate these results. We first conducted a qPCR experiment in which we grew THP1 cells as a mix of 14% perturbed cells (for either *NFKB1* or *TRAF6* and labeled with GFP in each case) with non-perturbed controls and seeded in different wells with two densities (low density: 25,000 cells/cm^2^, high density: 216,000 cells/ cm^2^). After LPS stimulus, we sorted perturbed and control cells from each density by FACS and extracted their RNA to quantify the expression levels of genes of *TNF*, *IL1a* and *ETV3*. Most differences of effect sizes are concordant between qPCR and Perturb-FISH, (**Fig 4c**) with the notable exception of the effect of *NFKB1* on *TNF*. The dependency of bulk *TNF* expression with average cell density has been reported before, albeit with inconsistent directions^27,28^. We therefore used RNAscope^31^ to better characterize this effect at the single-cell level since it is one of the perturbations most affected by density according to Perturb-FISH (LFC_low_: 0.45, Q-value: 0.015, LFC_high_: 0.007) We reasoned that an assay that does not require averaging expression across cells that are exposed to varied densities in a dish **(Supplementary Fig 1d)** would have better sensitivity. We find that RNAscope does confirm the Perturb-FISH findings: *NFKB1*-KO cells overexpress *TNF* at low density but repress it at high density (**Fig 4d**, and **Supplementary Fig 2**). We also used RNAscope to validate the finding that *IL1a* is expressed at lower levels in macrophages at high density (**Supplementary Fig 3a**) RNAscope data shows that *TRAF6* KO causes a complete silencing of *IL1a* regardless of local cell density (**Supplementary Fig 3b**). Therefore, the density-related difference in effects evaluated with Perturb-FISH originates from the density dependent difference in expression of *IL1a* in control cells, as this changes the value used as a reference in the comparison. Overall, we find that Perturb-FISH uncovers density related effects of genetic perturbations that are validated by other experimental approaches. These effects highlight the existence of mechanisms that regulate immunity at a cell population level and could be relevant to diseases related to over or under activation of the immune response and inflammation.

Next, we sought to identify intercellular effects of genetic perturbation on gene expression in neighboring cells. We analyzed cells that received a control guide and looked for effects coming from perturbed neighbors. We find 9 perturbations have significant effect on gene expression in their control neighbor cells, including *MYD88* (**Fig 4e**). Cells with a *MYD88* KO provoke an increase of *TNF* (LFC: 0.83, q-value: 0.06) and *CD14* (LFC: 0.77, q-value: 0.018) in their neighbor. *MAP2K7* perturbation also has several effects in neighboring cells, downregulating *TOP1* (LFC: - 0.75, q-value:0.06), *RPRD2* (LFC: −1.01, q-value: 0.028) and *CHD1* (LFC: −1.08, q-value: 0.06). While this analysis does not reach statistically significant effects from *IRF3* perturbation on *IL1a*, it suggests that *IL1a* is overexpressed in cells that neighbor an *IRF3* KO cell (LFC: 1.65, Q-value: 0.7) which is validated by our RNAscope data. (**Supplementary Fig 3**)

These results reveal coregulatory gene networks between macrophages responding to a bacterial infection and highlight the capacity of Perturb-FISH to interrogate genetic interactions between cells, which we believe will be a key tool in understanding how multicellular circuits underlie tissue function.

### Perturb-FISH for functional screens: interrogating autism spectrum disorder risk gene networks in disrupted calcium activity

To demonstrate the ability of Perturb-FISH to match perturbation data with functional imaging, we then applied it to study autism spectrum disorder (ASD) risk genes and their role in regulating astrocyte expression and activity. Whole exome sequencing studies have recently identified a large number of highly penetrant but rare risk genes associated with ASD but little is known about their molecular or phenotypic role, especially in the brain. A tempting hypothesis is that these risk genes converge on a smaller set of molecular and cellular pathways^32^. Several publications have highlighted transcriptional differences within astrocytes between ASD patients and control groups^33–35^. There have also been reports of a functional role of calcium activity in the development of ASD symptoms^36–38^, specifically that ASD patient iPS-derived astrocytes show aberrant calcium activity and that injecting these cells into healthy mice is enough to generate ASD like behaviors^39^; or that disrupting calcium activity in astrocytes generates ASD like behaviors in mice^38^. We thus decided to explore the power of Perturb-FISH to study a new biological problem by determining if a set of ASD risk genes with largely unknown roles converge on aberrant calcium signaling in iPS-derived astrocytes.

We designed a Perturb-FISH screen to determine if knocking down highly penetrant ASD risk genes^40^ also leads to dysfunction of calcium activity. We combined 50 ASD risk genes prioritized by the Simons Foundation Autism Research Initiative (SFARI), with additional risk genes identified in a whole exome sequencing study^40^ for a total of 127 risk genes and targeted these genes using CRISPRi. We perturbed these genes in astrocytes differentiated from the UCSFi001-A line (human pluripotent stem cells from healthy male donor), stably expressing dCas9-KRAB and previously used for CRISPRi based expression and morphology screens^41^. Since calcium transients have been well characterized to follow ATP stimulation in astrocytes^42–44^, and since astrocyte intrinsic calcium activity following ATP stimulation is a result of *ITPR* mediated release of calcium ions from the endoplasmic reticulum (ER), we also included gRNAs against *ITPR3* as positive controls^45^. The genes selected for expression measurements included the perturbed targets, and a published list of 358 differentially expressed genes between astrocytes from ASD and control patients^34^.

To characterize functional profiles, we stimulated astrocytes with 250 μM ATP, and recorded calcium activity in 26,000 cells (**Fig 2d**). We parametrized the calcium traces for every cell (**Methods**) to identify cells that responded to the stimulation by starting calcium oscillations. As shown previously by our group and others^46,47^ this parameterization allows clustering of astrocytes in several groups with or without transients of several profiles, where a transient is a brief increase of calcium concentration in the cytoplasm: “inactive cells”, “with transients, high plateau”, “no transients, high plateau”, “large transients”, “no transients, low plateau”, and finally “delayed transients. (**Fig 5a** and **Supplementary Fig 4a**). Notably, the cluster of cells “with transients, high plateau” best resembles the activity pattern found in ASD astrocyte lines in a previously reported ASD case-control cohort^39^.

**Figure 5.**
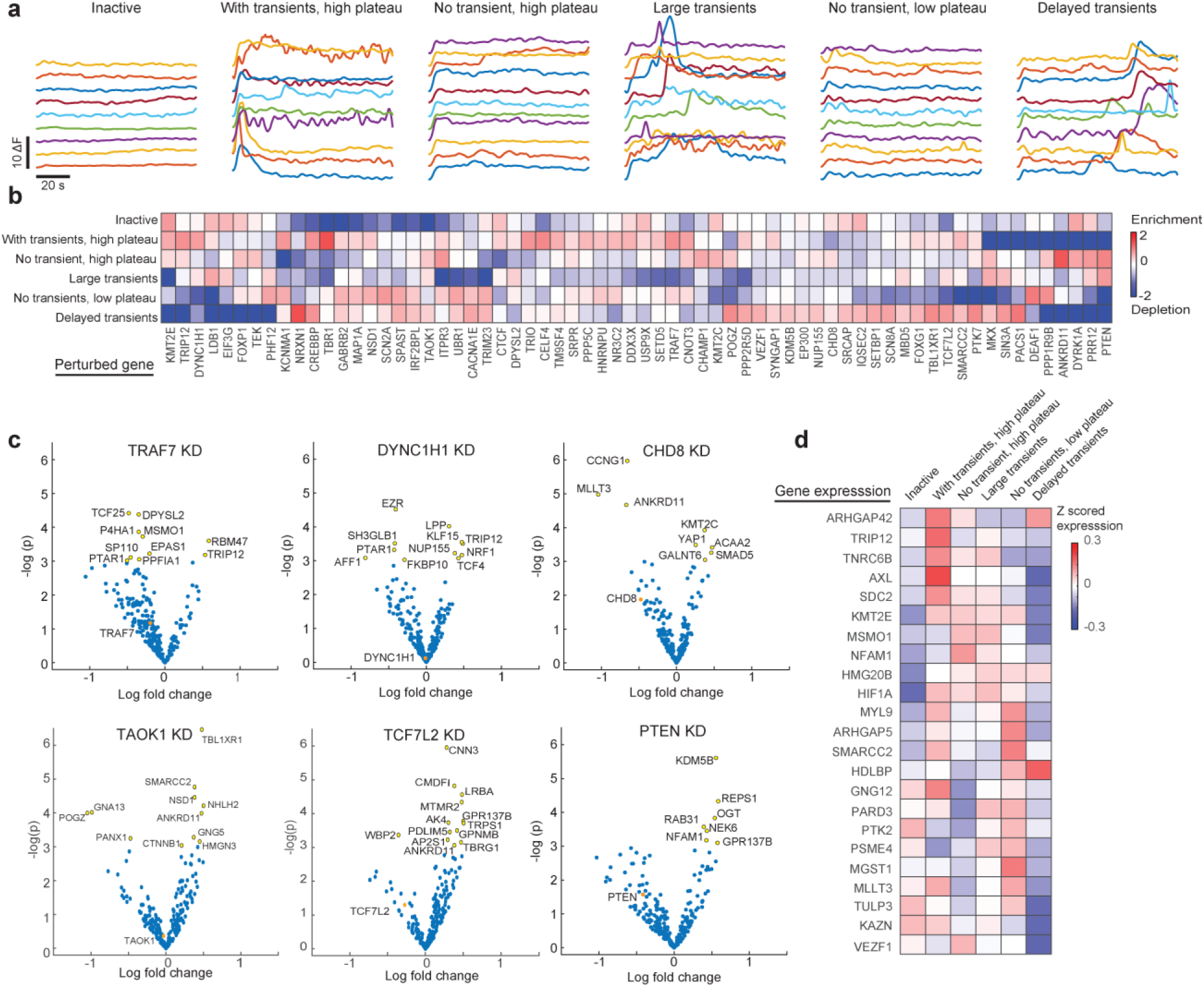
Perturb-FISH analysis of the role of ASD risk genes in generating different calcium activity phenotypes in astrocytes. A) Traces showing the variation in cytoplasmic calcium (y-axis) concentration in example cells (line) across time (x-axis), following ATP stimulation. Variation in calcium concentration shown as difference in fluorescence. Cells are 6 groups using k-means clustering on features extracted from the traces: inactive cells, cells with transients and a high plateau, cells without transients but with a high plateau, cells with large transients, cells without transients and with a low plateau, and finally cells with delayed transients. B) Heatmap showing the enrichment (scale bar) of perturbations in 6 clusters of calcium activity, measured as the ratio between detected and expected frequency, and shown as log fold change in natural log base. C) Volcano plots showing LFC effect sizes (x-axis) versus significance (y-axis) for 6 example perturbations. Yellow indicate significantly affected genes (p<0.05), orange indicates the effect of the perturbation on its target gene. D) Heatmap showing genes with differential expression (x-axis) and their z-scored expression (scale bar) in 6 clusters of calcium activity.

Coupling Perturb-FISH to a functional readout allowed us to perform a gRNA enrichment analysis akin to optical pooled screening^10^. We quantified gRNA enrichment per cluster (**Fig 5b**) and a p-value of being enriched (**Supplementary Fig 4**). Overall, all calcium response patterns are present in control astrocytes, matching our previous knowledge of the system^48–50^, but perturbations to certain ASD risk genes push cells to specific functional profiles. Unsurprisingly, *ITPR3* KD cells do not generally have transients and are highly enriched in the cluster of cells “no transients, high plateau” (LFC: 0.8, p-value: 0.004). Consistent with the idea that ASD risk genes can have common phenotypic effects, we found groups of significant perturbations enriched in each functional cluster: *DYRK1A*, *CTCF, KMT2E* and *KMT2C* in inactive cells, *ANKRD11*, *TRIP12*, *CHAMP1*, *CNOT3* in cells with “no transients, high plateau”, *DEAF1*, *NSD1*, *SCN2A* in cells with “no transients, small plateau”, *LDB1*, *PTEN* and *SIN3A* in cells with “large transients” and *NRXN1*, *PACS1* in cells with “delayed transients”. Interestingly, a large number of ASD risk genes perturbations were significantly enriched in the “with transients, high plateau” cluster (*CECL4*, *CNOT3*, *CREBBP*, *KCNMA1, TBR1*, *TRAF7*, *TRIP12*) offering a hint to the potential cause of previous reports showing increased, noisier activation of calcium release following ATP stimulation in ASD derived astrocytes^39^.

The simultaneous mRNA expression readout of Perturb-FISH allows us to go beyond gRNA enrichment to nominate molecular mechanisms underlying the observed functional phenotypes. In the absence of prior knowledge of the system, we applied stringent filters on the expression data. We identified lowly expressed genes and removed both perturbation and expression level information for these genes from the dataset. We also removed all perturbations present in fewer than 10 cells leaving 78 perturbations and 237 measured genes. We then divided the data into two halves and showed that significant effects inferred from each half are 68% correlated. (**Supplementary Fig 4b**) After computing the effects on this filtered dataset, we find 798 effects with a nominally-significant p-value below 0.05, of which 44 have an absolute value larger than 1 (meaning a difference in raw counts at least 2.7 fold), impacting 35 genes and caused by 26 perturbations. (**Supplementary Fig 5**).

We jointly analyzed transcriptional effects with calcium profile enrichment data to formulate hypotheses on the mechanism of these enrichments. We first examined *TRAF7* KD cells, which were over-represented in the cluster “with transients, high plateau”. *TRAF7* KD led to a downregulation of *DPYSL2* (LFC: −0.35, p: 0.01) and an upregulation of *RMB47* (LFC: 0.58, p: 0.02) and *TRIP12* (LFC:0.53, p: 0.04) (**Fig 5c**). *DPYSL2* has been reported to promote a neuronal lineage over an astrocytic lineage^51^; while *RBM47* was also found to promote a neuroectoderm lineage^52^; and *TRIP12* was shown to repress *FBW7*^53^, a gene that was shown to push neural progenitors to differentiate into astrocytes^54,55^. We also observed that *DYNC1H1* KD, another gene over-represented in the “with transients, high plateau”, led to overexpression of *TRIP12* (LFC: 0.49, p: 0.03) (**Fig 5c**). Combined, these results suggest a link between genes promoting an astrocyte lineage from neuronal progenitors and astrocytes more responsive to ATP stimulation.

We found interesting connections between ASD risk genes which did not correlate with calcium phenotypes. For example, *ANKRD11* is affected by several perturbations: *CHD8* knock down (**Figure 5c**) causes its downregulation (LFC: −0.66, p-value: 0.009), while knocking down *TCF7L2* (LFC: 0.38, p-value:0.046) or *TAOK1* (LFC; 0.47, p-value: 0.018) causes an overexpression of *ANKRD11* (**Supplementary table 1**). *CHD8* has previously been reported to bind *ANKRD11* at the protein level, so this data suggests an additional layer of regulation at the transcript level, as well as an interplay with several other ASD risk genes. However, given that these genes all have different functional signatures, this suggests that any potential connections are not related to the calcium phenotypes observed.

To explore this further, we use the phenotypic clusters to reduce the dimensionality of the data: pooling together cells that share a phenotype increased statistical power when comparing transcriptomic signals. We examined the gene expression signature of each functional cluster, finding 23 genes that are differentially expressed. (**Figure 5d** and **Supplementary table 2**), with each functional cluster characterized by different expression profiles. (**Fig 5d**). Focusing again on the cluster “with transients, high plateau”, we found that *TRIP12* is strongly up-regulated (Zscored expression: 0.17, p-value: 0.04), as are *ARHGAP42* (Zscore: 0.15, p-value: 0.006), *TNRC6B* (Zscore: 0.13, p-value 0.007) and *AXL* (Zscore: 0.21, p-value 0.0007). *AXL* has been linked to phagocytosis in activated astrocytes^56^. *ARHGAP42* is one of the genes that is most downregulated following general anesthesia, an effect that is thought to be a response to the lower local activity of the brain^57^. *TNRC6b* was linked to reduced frequency but not amplitude of excitatory currents in response to depletion of *CPEB4*^58^. Together, these data establish a connection between several high-confidence ASD risk genes and increased activity in astrocytes. We elaborate below (**Discussion**) on this connection and potential roles in disruption of neuronal activity.

We also describe the example of two perturbations (*PTEN* and *TRAF7*) with connected effects across all three data modalities, making them good candidates for further characterization. First, cells knocked down for *PTEN* are enriched in both the “large transients” and the “no transients, high plateau” clusters (LFC: 0.78, p-value: 0.05 and LFC: 0.94, p-value: 0.047 respectively; making up almost all cells with a *PTEN* gRNA) (**Fig 5c**). The knock down of *PTEN* causes an upregulation of the expression of *NFAM1* (**Fig 5b**). *NFAM1* is indeed overexpressed in the cluster “No transients, high plateau” (Zscore: 0.12, p-value: 0.01 **Fig 5d**). *NFAM1* is a transmembrane receptor with a documented role in hyperactivation of immune pathways^59^, but its role in ASD pathology is not understood^60^. Our second example is about *TRAF7*: cells knocked down for *TRAF7* are enriched (LFC: 0.85, p-value: 0.01) in the cluster “Large step with transients”. (Fig 5c) *TRAF7* KD results in an overexpression of *TRIP12* (LFC: 0.53, p-value: 0.04, Fig 5b), which is indeed overexpressed in that cluster (Zscore: 0.17, p-value: 0.04, Fig 5d)) Such connections suggest that *TRAF7* is regulating the level of expression of *TRIP12,* and suppressing it leads to hyper-active cells through the overexpression of *TRIP12*. These results demonstrate how our data can be used as a powerful tool to screen not only gene interaction networks within and between cells, but also as a new screening approach for the functional roles of genes in cell systems.

## Discussion

In the present study, we developed and evaluated a new protocol, Perturb-FISH, using all-optical experimental methods to study the genetic and molecular associations of spatial and functional biology at single-cell resolution. Our analysis demonstrates that Perturb-FISH results produce robust inference of intracellular perturbation effects with power at least as good as Perturb-Seq, with the caveat (discussed below) that Perturb-FISH captures expression for only a targeted set of genes. We further present a series of results that cannot be found using conventional Perturb-seq, as they depend on the spatial context or functional profiling of each cell.

Intracellular effects in our first screen (macrophages responding to LPS stimulus) have been well-characterized, but using Perturb-FISH allowed us to observe how these responses depend on spatial context. Notably, we found that several hallmark genes of the innate immune response are differentially regulated depending on local cell density, and we observed that expression in control cells can be affected by perturbations in neighboring cells. Such density-dependent and intercellular effects at a single-cell level likely have a profound effect on population-level immune response and the balance between pro- and anti-inflammatory factors, with further studies needed to clarify how the environment shapes cellular responses and how cellular activity (or perturbation) shapes local environment.

A critical advantage of Perturb-FISH is the ability to match the intracellular effects of genetic perturbations with a functional readout, such as live calcium imaging. We demonstrated the feasibility of the experiment by screening ASD risk genes in astrocytes and recording calcium activity in response to an ATP stimulation. Unlike the THP1 screen, relatively little is known about the role of the newly identified risk genes. By linking genetic perturbation with gene expression and functional profiles we were able to generate hypotheses about several of these risk genes regarding their effects on the transcriptome and role in calcium activity both in the form of genes that are differentially expressed in cells of different phenotypes, but also as targets whose silencing sets a particular phenotype for the cell. In particular, we find several targets that push the cells to favor one of the naturally occurring calcium phenotypes. This may be of special interest as previous works have reported hyper calcium activity in ASD patients and cells from ASD patients^39^ which results in a disruption of neuronal activity by impairing the controlled development of a synapse network. Our Perturb-FISH screen found several ASD risk genes which yielded similar functional phenotypes, and are thus candidates to explain the observed behavior in ASD patient lines. While establishing simple regulatory relations between disease risk genes will not be enough to understand how their dysregulation generated a pathology, Perturb-FISH can be used to find genes that act on similar pathways and to narrow the number of low throughput experiments necessary to elucidate the mechanism of action of risk genes. When used in conjunction with functional profiling, Perturb-FISH is not just a screen of gene interactions but also a screen for gene function.

With a guide barcoding strategy that mimics MERFISH, Perturb-FISH offers the same scalability in number of perturbed genes as MERFISH does in terms of recorded genes. For instance, as many as 140 gRNAs can be identified in 15 images (bits) with 5 rounds of imaging in 3 colors, or as many as 316 gRNAs in 21 bits, or 550 in 25 bits. The number of genes that MERFISH records scales identically with the number of cycles, and the number of imaging cycles needed for Perturb-FISH is the sum of both. Furthermore, we have shown that CRISPR-KO and CRISPR-i are both compatible with Perturb-FISH, but, in principle, the method is compatible with any of the array of gRNA-based perturbations including base editing, CRISRP-a, CRISPR-off, and non Cas9 based systems such as Cas12a or Cas13.

A key limitation of our method is the focus on a targeted set of genes for transcriptomic readout, which limits the potential for *de novo* discovery. This limitation can be addressed in the future in several ways. First, while we have designed the chemistry around MERFISH binding and readout, we note that multiple hybridization-based chemistries are compatible with Perturb-FISH, including CosMx^61^, STARmap^62^, and Seq-FISH+^63^. Notably, each of these has been shown to be compatible with higher plexes, up to 10,000 genes. Second, while we relied on commercial reagents to simplify development, we note that a fully homebrew approach could decrease reagents costs dramatically (as much 10-fold) and thus increase the number of genes profiled and perturbed. Finally, optical improvements maximizing the coverslip active area can substantially increase the number of cells profiled per dollar. Thus, we believe that Perturb-FISH is best thought of not as an alternative to scRNA-seq, but as a separate tool used to profile phenotypes dependent on a cell’s native environment and guided by alternative experiments (*e.g.,* our knowledge of the LPS response pathway or of ASD-involved genes).

The ability to interrogate the extracellular effects of cellular perturbations is a key step in understanding how they might generate function at the tissue scale. Using Perturb-FISH to decipher intercellular networks will require thinking differently about CRISPR screens. One might generally want to scale up the experiment to screen as many genes as possible; and Perturb-FISH can indeed achieve large numbers of perturbed genes, but looking at their effects on the neighbor cells require finding enough occurrences of a particular cell pair (one control touching one perturbed cell). The frequency of such occurrences is the product of the proportion of controls and a specific perturbation. Therefore, if an experiment is designed to measure the extrinsic effects of guide perturbations, the relative proportion of control guides will directly affect the power of the screen (as will total number of cells, cell density, and number of perturbations), and hence such screens will likely need to include a much higher proportion of control guides than a typical Perturb-Seq screen.

Overall, we believe Perturb-FISH advances the scale of an emerging foundational method in functional genomics for the study of intracellular circuitry, while also enabling the study of the genetics of intercellular circuitry with rich molecular detail.

## Acknowledgments

We thank Martin Kampmann for providing the line of astrocytes used in this work. This work was supported by NIH grants R01MH128366 and RF1MH121289, and by the Israel Science Foundation.

## Author contributions

Conceptualization, L.B., V.V., S.F. and B.C., Investigation, L.B., S.D., J.B., B.S., writing-original draft, L.B., S.F, B.C., writing-review and editing, L.B., V.V, R.N., S.F, B.C., software and formal analysis L.B., S.F., and B.C., supervision and funding acquisition R.N., S.F. and B.C.

## Methods

### Selecting genes to be measured with MERFISH

We used data from our matched Perturb-Seq^1^ screen to select an informative list of 130 genes to measure with MERFISH in our LPS response screen. Perturb-Seq data allowed us to establish modules of genes with correlated expression levels after LPS-stimulation. We selected a list of genes capturing major directions of variation (as determined by PCA), with a total expression across these genes below 2000 FPKM, and no individual gene above 30 FPKM which are conditions imposed by the use of MERFISH: high abundance creates crowding in the image and prevents decoding. In brief, we ran a principal component analysis on the raw counts matrix, and scored every gene for how much of the variance of a principal component they captured. The list of genes was built by iteratively adding new genes. At each iteration, we simply cropped the original count matrix to estimate the count table we would get if we ran MERFISH with that list of genes. We then used a linear regression to fit this estimated data onto the PCA space, and calculated residuals and explained variance. The principal component with the lowest explained variance received 2 additional genes, which were chosen for being the most correlated with the residual.

For the ASD screen, we used a list of risk genes published by the SFARI consortium (50 genes), to which we added genes from a published work for Sattertrom^64^ and control genes known to regulate ATP signaling of calcium activity, for a total of 127 target genes. Since little is known about most of the 127 genes selected for perturbation, we chose to record the same list of genes with MERFISH. In addition, we selected 358 genes that are differentially expressed in astrocytes between healthy patients and people with ASD^65,66^, totaling 485 genes.

### Virus preparation

Viruses were prepared using a previously published protocol (https://portals.broadinstitute.org/gpp/public/dir/download?dirpath=protocols/production&filename=TRC%20shRNA%20sgRNA%20ORF%20Low%20Throughput%20Viral%20Production%20201506.pdf). Cas9 virus was concentrated by centrifugation through a 100kD purification column.

### Generating Cas9 expressing cell lines

THP1 cells from ATCC® (TIB-202™) were maintained in RPMI (ATCC, RPMI-1640) with 10% FBS (ATCC® 30-2020) without antibiotics. Cells were split to maintain a density between 500k cell/mL and 1.8M cells/mL. THP1s were infected with Cas9 virus by preparing 20 wells of a 96 well plate with 200k cells per well, with 50μL medium and 150μL virus (in medium) in 8μg/mL polybrene. Cells were spinfected at 1000g for 90 min, then placed overnight in an incubator at 37C. The following morning medium was replaced with fresh maintenance medium and cells were left to recover for 2 days. Cells were then selected in medium containing 6μg/mL basticidin until control wells that received no virus were dead (8 days). Cells were then amplified for 20 days and stocks were frozen. We assessed Cas9 activity by infecting these cells (same protocol) with a virus containing both GFP and a guide against GFP. After puromycin selection, we used cytometry to quantify the number of cells that were not GFP positive (cells in which GFP was knocked out by Cas9 activity) and found 73% activity (**Supplementary Fig 6**).

Astrocytes were differentiated from dCas9-KRAB expressing hi-PSCs following our previously published protocol^48^.

### Vector design and cloning guide libraries

LentiGuide-Puro was a gift from Feng Zhang (Addgene plasmid # 52963). The vector was modified to replace the sequence of the gRNA scaffold with the optimized sequence from Hill et al. ^67^, add a T7 between the end of the U6 and the cut site of BsmbI^20^, we also removed the T7 region that is part of the LentiGuide-Puro to only keep one. The final plasmid was purchased from Genscript.

We designed libraries containing 2 guide RNAs per target gene. Guides were designed using the CRISPick tool from the Broad institute 30 base pair cloning Gibson arms were added on either side during synthesis. These are complementary to the flanking regions on the vector plasmid and are therefore the T7 promoter on the 5’ side, and part of the optimized scaffold on the 3’ side. We manually replaced guides (with the next best guide from CRISPick^68,69^) when stitching these regions resulted in sequences of more than 3 consecutive Gs as we found these impaired proper amplification before cloning and presumably also during T7 transcription. Guide libraries were ordered as oPools from IDT (libraries were ordered several times within one single oPool to minimize variations in guide frequencies). The library was PCR amplified and assembled using a Gibson assembly. The transformed plasmids were electroporated into competent bacteria (electroporation results in more uniform libraries than heat shock) and maxi prepped. 2 non targeting guides (not matching anywhere in the genome), and 2 safe targeting guides (matching a non-coding region) were cloned separately and added at the end to represent 30% of the final library. The final library was sequenced using a MiSeq to verify guide distribution. Briefly, two PCRs were used to amplify the guide region, and first add the proper read1 and read2 priming regions then the P5 and P7 regions. We spiked 30% PhiX into the library before loading on the MiSeq to compensate the relative low variability of this library.

### Infecting cells with guides

Cas9 THP1 cells were infected with guides by spinfecting 40 wells of a 96 well plate with 200k cells per well, with 2μL of virus and 8μg/mL polybrene for one hour at 800g. Spinfection was done in the evening, and the following morning the cells were washed and placed in normal medium at 800k cells /mL. After one day rest, cells were selected in puromycin 4μg/mL until a control well was fully dead (2 to 3 days). The amount of guide virus to use was established by infecting cells with serial amounts of virus (from 0.5μL to 50μL), selecting cells with 4μg/mL puromycin for 3 days. After selection, numbers of cells were compared to a well that was spun with polybrene but no virus to find the amount of virus to use to obtain 25% survival, which corresponds to a MOI (multiplicity of infection) of 0.25.

Differentiation of human astrocytes from i-PSCs was conducted for 30 days, following our previously published protocol^5^ Cells were maintained in Astrocyte Medium (ScienCell, Cat #1801), and were infected with the guide library at day 24, to be ready for calcium imaging and Perturb-FISH at day 30.

### THP1 plating and stimulation

THP1 cells were plated in a polyornithine/laminin coated 6 well plate, at 1 million cells per well, in medium containing 20ng/mL phorbol-12-myristate-13-acetate (PMA). After 24 hours, medium was replaced with fresh medium without PMA. 2 hours later, cells were lifted with Trypsin and replated on polyornithine/laminin glass. The glass used was the circular coverslip compatible with MERFISH, provided by Vizgen. We used removable chambers from IBIDI to restrict the plating area and only plate the number of cells that could be later imaged. 42k cells were plated in 0.22 cm² for the negative control, and 100,000 cells were seeded in 0.56 cm² chambers, on the same glass slide, to generate wells that would be later stimulated with LPS. One day later, medium was replaced with either fresh medium or medium containing 100ng/mL LPS. Cells were then incubated for 3 hours prior to fixation.

### Guide RNA codebook design

Following the approach used in MERFISH, we use a Hamming weight 4 Hamming distance 4 codebook to encode the identities of our guide RNAs. Every guide is encoded as a binary barcode that contains exactly 4 “ones” and any 2 barcodes from the codebook are different on a least 4 bits. This approach allows for single bit error correction as a barcode that contains an error has only one bit different from its actual matching code, and at least 3 bits difference with all others. Barcodes that contain 2 errors are discarded as they could match 2 possible barcodes. We find that certain combinations of ones and zeros are particularly sensitive to false positive detection, or dropout and should be avoided: all sequences that would result in a guide always lighting up in the same color, or in a guide being “on” in all 3 images in one single round should be avoided.

### Perturb-FISH encoding probes

After T7 transcription, each cell contains multiple copies of RNA at the site of genomic insertion. These RNA molecules started with the gRNA sequence followed by the optimized guide scaffold used in our vector. We targeted these regions with encoding probes that contained a 30 base pair annealing region matching the gRNA and the first 10 base pairs (bp) of the optimized scaffold. This allowed a hybridization region of 30 base pairs, which is also the length used in the MERFISH probes, therefore allowing similar sensitivities to temperatures and formamide concentrations. Each hybridization region was flanked with two 20 bp readout regions on either side that were complementary to the 4 readout probes used to encode the 4 bits that characterized each gRNA.

### Perturb-FISH readout probes

For the THP1 experiment, all readouts (for guides and genes) were purchased as a kit from Vizgen. For the astrocytes, fluorescent readout probes were acquired from Eurofins with sequences matching those published by C. Xia et al.^70^ (Supplementary Table 1). These oligos are tagged with a fluorophore attached to the 5’ side with a disulfide bond. The readout buffer was made of 10mL 50% ethylene carbonate 5mL 20X SSC, 34 mL RNAse free water, 1mL 25% Triton-X 100, 200uL tween-20 and 50μL murine RNAseI as used before by others^70^.

### Perturb-FISH protocol

Cells were washed twice in PBS, then fixed on ice in methanol/acetic acid at a 3:1 ratio as done by A. Askary et al^17^. As the authors of this work note, it is important not to fix with PFA as this blocks T7 transcription. Cells were washed in PBS by doing exchanges to prevent the sample from completely drying. Coverslips were placed in a 60cm petri dish, filled with PBS, and the IBIDI chambers were then peeled from the glass, allowing all conditions to receive the same later incubations. T7 transcription mix was prepared following manufacturer (Thermofisher, AM1334) instructions: 8μL of each dNTP was mixed with 8uL 10x reaction buffer, 8μL T7 enzyme, and 32μL nuclease free water. The liquid covering the coverslip was then removed, a drop of the transcription mix was pipetted on top of the areas with cells, and covered with a square of parafilm to spread the drop and prevent evaporation. Coverslips were incubated at 37C for 6 hours, then fixed without washing, in 4% PFA, for 15 minutes, then washed 3 times 5 minutes in PBS and stored overnight in 70% ethanol. Coverslips then underwent normal MERFISH preparation: samples were washed in formamide wash (5mL 20X SSC, 15 mL 100% deionized formamide, 30mL nuclease free water) at 37C for 30 minutes. An encoding buffer was then prepared by mixing the MERFISH encoding library provided by VIZGEN with probes targeting the guide RNA regions from a stock at 300X concentration to reach a final concentration of 1nM per probe, plus 4nM of 5’acrydite-modified guide anchors GCGCCAAAGTGGATCTCTGCTGTCCCTGTAA, and GGATGAATACTGCCATTTGTCTCAAGATCTA that anneal to the T7 transcribed RNA downstream of the scaffold, and will later anchor the transcripts to the acrylamide gel. The coverslips were placed overnight at 37C in a cell culture incubator. The sample was then washed twice for 30 minutes in formamide wash buffer at 47C. Finally, the sample was washed 3 times in 2X SSC for 2 minutes each, then incubated for 10 minutes in 5mL 2X SSC containing 1μL of 5000X fiducial beads solution (Polysciences, #17149-10). The sample was then embedded in a polyacrylamide gel, then cleared for one day at 37C following the protocol used in MERFISH. When necessary, the samples were stored in 2X SSC containing 1μL/mL murine RNAseI. After clearing, samples were bleached in 2X SSC for up to 12 hours under a blue LED to further reduce background noise.

### Sample Imaging

Cells were stained with DAPI 1/1000 for 5 minutes and imaged on an alpha version of Vizgen’s MERSCOPE. Images dedicated to the detection of guide RNAs were acquired first. Briefly, 5 automated cycles of imaging were run to collect all 15 bits encoding the guide library used on THP1s. Each cycle started with 15 minutes incubation of the next readout buffer (containing 3 readout probes with dyes at 750nm, 647nm, and 565nm). The samples were then washed for 15 minutes in wash buffer (5mL 20X SSC, 10mL 50% ethylene carbonate, 35mL RNAse free water). The wash buffer was then replaced with imaging buffer made of 15mL 2X SSC, 1.66mL Trolox quinone at 500μM [Mix 395mg Trolox in 15.8mL methanol. Dilute 0.4mL of that solution in 20mL 2X SSC. Place in UV crosslinking oven and bake for 1 hour at 254 nm. Measure absorbance at 255. The concentration in μM is (abs(255nm) – 0.8)/11200*1000000. Dilute in 2X SSC to 500 μM], 33μL protocatechuic acid (2.5M), 83μL Trolox 10% in methanol, 33μL recombinant protocatechuate 3,4-dioxygenase, 33μL RNAseI, and NaOH to adjust pH to 7. Each imaging round finished with a 15 minutes incubation in extinguishing buffer (2X SSC with 50mM TCEP) followed by a 10 minutes rinsing step in 2X SSC with RNAseI. After acquisition of the images of the guide RNAs, the transcriptome was imaged using reagents provided by VIZGEN. It is critical that buffers do not stay at room temperature for too long as their degradation impairs the detection of transcripts/guides, and we therefore ensured that the automated fluidics system was only loaded with the buffers required for the next 12 hours of imaging. Buffers were added to the fluidics system as imaging progressed as a complete Perturb-FISH run can easily require 5 days of imaging.

### Decoding guide identity

Raw images were registered using previously published algorithms used for MERFISH image analysis^71^. Image preprocessing was performed similarly to Lu et al^72^. Images were divided by images of the background in each channel to correct for uneven illumination across field of views. Spots were then enhanced with a gaussian filter of size 2 using MATLAB’s imgaussfilt. Background was computed as the same image filtered with a gaussian filter of size 12, and subtracted. Image stacks of 7 Z-levels were then max-projected. These images were then binarized using a simple threshold, and filtered to remove spots smaller than 7 pixels or bigger than 450 or with an eccentricity over 0.8. For each field of view, images from all channels were then max-projected to find the locations of putative gRNAs. For each of these putative guides, a centroid position was determined, and the sequence of images that contained a binary object within a radius of 5 pixel of this centroid was compared to the codebook of the guide library. The use of a Hamming weight 4 Hamming distance 4^13,14^ codebook allows identification of all guides that fluoresce in 3, 4, or 5 images. We then used the standard deviation of spot intensity and their average fluorescence across image rounds to generate scatter plots (Supplementary Fig 6). On these plots, 2 clouds of points were easily identifiable, one of them is enriched in blank barcodes than do not actually match any gRNA from the library. We therefore considered this cloud as misidentified gRNAs and filtered these false gRNAs out of our analysis.

mRNA images were decoded using MERlin with default parameters^71^. (https://github.com/emanuega/MERlin)

### Cell segmentation

THP1 cells were segmented using a watershed algorithm. Nuclei were segmented on each field of view, and a binary mask of nuclei centroids was then created to use as seeds for the watershed algorithm. To generate basins, we used a map of all transcripts decoded by running Merlin on our MERFISH images, then used a gaussian filter to blur them out, resulting in a mRNA density map of the sample.

Astrocyte nuclei were segmented using a binary mask of DAPI signal, then morphologically dilated this mask. These images (acquired at 60X) were downscaled to match the dimension of the calcium images (2X) and recognizable features in the dish were manually identified and used to compute an affine geometrical transformation using MATLAB’s *estimateGeometricTransform* function to realign images across modalities. Images were then translated using *imwarp_same*.

### Generating count tables

Count tables were assembled using previously built masks and matrices of decoded transcripts: for each object of the mask, the number of transcripts for each gene was summed. Design matrices containing the identity of the perturbation received by each cell were assembled in the same way. THP1 expression tables were filtered to remove cells with less than 70 or more than 2300 counts. Expression tables for the astrocytes were filtered to keep cells with counts between 200 and 1900, and genes with at least 0.5 counts per cell.

### Determining perturbation effect sizes

The effects of perturbations in THP1s were determined using our published tool FR-Perturb^1^, using a rank of 34, lambda1 and lambda2 of 0, and log_exp_baseline of 2.7 and the total number of transcripts per cell as a covariate. The script was slightly modified to remove the normalization to 10,000 counts per cell that is traditionally performed on droplet-based data. These parameters were determined by screening a range of values and keeping the ones that resulted in the highest correlation between effects computed on two random halves of the data. FR-Perturb outputs two tables: one contains the effects of the perturbations expressed as log fold change, the other contains q values for significance analysis. Significant effects are here defined as effects with associated q value below 0.1.

In the part of the study in which we consider density, the population of cells was first divided depending on the number of neighbors of each cell. Low density was defined as 2 or fewer neighbors, and high density as 3 or more.

In the case of astrocytes, perturbed target on gene expression effects were computed by simply averaging expression counts in all cells that received a given perturbation, and dividing these by the average counts in control cells. For each gene, significance was evaluated with a Student’s T-test between perturbed cells and control cells.

Log-fold-change (LFC) values all use natural log, all correlations are Pearson’s correlation.

### Neighbor detection

To identify the neighbors of a given cell, we dilated its mask by 50 pixels (5.4um) in all directions in the MERFISH data, or by 10 pixels (6.2μm) in the RNAscope data. Any cell that overlapped with this dilated mask was identified as a neighbor. Cells with 2 or fewer neighbors were called “low density” cells, while cells with 3 or more neighbors were called “high density” cells.

### qPCR validation of effects

THP1 cells were transduced with a modified gRNA vector containing both puromycin resistance and GFP. After selection, cells were mixed at ratios of 14% perturbed cells with 86% non-perturbed controls (received a non-targeting guide), and seeded in 6 well plates with 20ng/mL PMA for differentiation. After a day, cells were re-seeded in fresh medium in a 48 well plate at different densities: low density: 25,000 cells/cm2, high density: 216,000 cells/ cm2. Cells were stimulated for three hours in LPS at 100ng/mL, then detached and sorted perturb and control cells for each density by FACS based on GFP expression, and extracted their RNA using a miniprep kit (NEB, T2010S), performed reverse transcription (NEB, E3010L) and ran qPCR (NEB, M3003L) in triplicates. Actin-B was used as a reference for transcript abundance.

### RNAscope data generation

For effect validation using RNAscope data, cells were transduced with a modified gRNA vector containing both puromycin resistance and GFP. Cells were prepared following the same protocol as for the qPCR and seeded in 96 well plates at the same density as before. Samples were stained following the manufacturer’s protocol^16^ and imaged at 10X on a Nikon Ti2-E microscope. GFP signal is used to identify cells that received a perturbation.

### Calcium imaging

Astrocytes were grown on glass coverslips (VIZGEN 10500001), in chambers made of PDMS (poly dimethyl sulfoxide). Cells were loaded for 30 minutes with 2uM Fura-4AM dye at 37 °C, then washed and imaged in recording solution (125mM NaCl, 2.5mM KCl, 15mM HEPES, 30mM glucose, 1mM MgCl_2_, 3mM CaCl_2_ in water, pH 7.3, mOsm 305) as detailed in M Berryer et al.^5^ Time lapse videos were acquired at 2X on a Nikon Ti2_E microscope equipped with a Photometrics Iris 9 camera at 2 Hz images per second under 488 nm illumination provided by a Lumencor Celesta, emission signal was acquired through a Chroma ET560/40x filter. Cells were stimulated by adding one volume of recording solution with 200μL of ATP at 500μM to the one volume already in the well, for a final concentration of 250μM. Cells were then fixed and carried through to the Perturb-FISH protocol.

For each cell, signal intensity across time was computed as the mean value of all pixels in the cell, using the aligned cell mask from MERFISH. The base level of each trace was subtracted, and peaks were detected using a *findpeak* algorithm from MATLAB, with thresholds set to 15 for the minimal height, 6 for the prominence (local height) and 15 for the space between peaks. Traces were not further normalized because the absolute height of the calcium response carried biological information.

For parametrization of this data, we used the number of peaks, the local peak height (prominence), the size of the first step (defined as the difference between the maximum value of calcium intensity in the 5th to 15th frames and the minimal value between the 1st and 10th frames), the first response time (*i.e.*, the delay before detection of the first peak, set to 200 – the duration of the analyzed video – when no peak was detected), the average peak delay, the area under the curve, and a binary parameter set to 1 if at least a transient was detected, 0 otherwise. These numbers were then Z-score normalized, and k-means clustering was used to cluster the cells into 6 groups of different signature calcium activity.

### Astrocyte differential calcium profile and expression analysis

We used the distribution of control cells across calcium trace clusters to establish an expected distribution of cells between clusters. For perturbation to a target “A” we calculate enrichment or depletion in a cluster “B” as the log-ratio of the proportion of “A” cells falling in cluster “B” with the expected frequency based on the control distribution. For statistical analysis, we used a hypergeometric test on the raw values of guide abundance, where the p-value for the enrichment of guide A in cluster B is 1-hygecdf(x-1,M,K,N) with hygecdf MATLAB’s hypergeometric cumulative distribution function, x the number of cells with guide A in cluster B, M the total number of cells, K the number of cells in cluster B, N the total number of cells with guide A.

Gene expression signatures for each cluster were computed as the mean of the Z-scored expression of a gene for all cells in a cluster. Significance was established using an ANOVA test on raw expression levels.

## Data availability

Image data, and tables of raw counts, perturbation design and calcium intensity will be made available at the time of final publication on SSPsyGene DRACC. Data from the matched Perturb-seq screen were deposited in the National Center for Biotechnology Information’s Gene Expression Omnibus under accession number GSE221321.

The vector will be available on addgene prior to final publication.

## Code availability

FR-Perturb is available in a public repository: https://github.com/douglasyao/FR-Perturb. The code used for image analysis will be made available on github prior to final publication.

## Supplementary Figures

**Supplementary figure 1.**
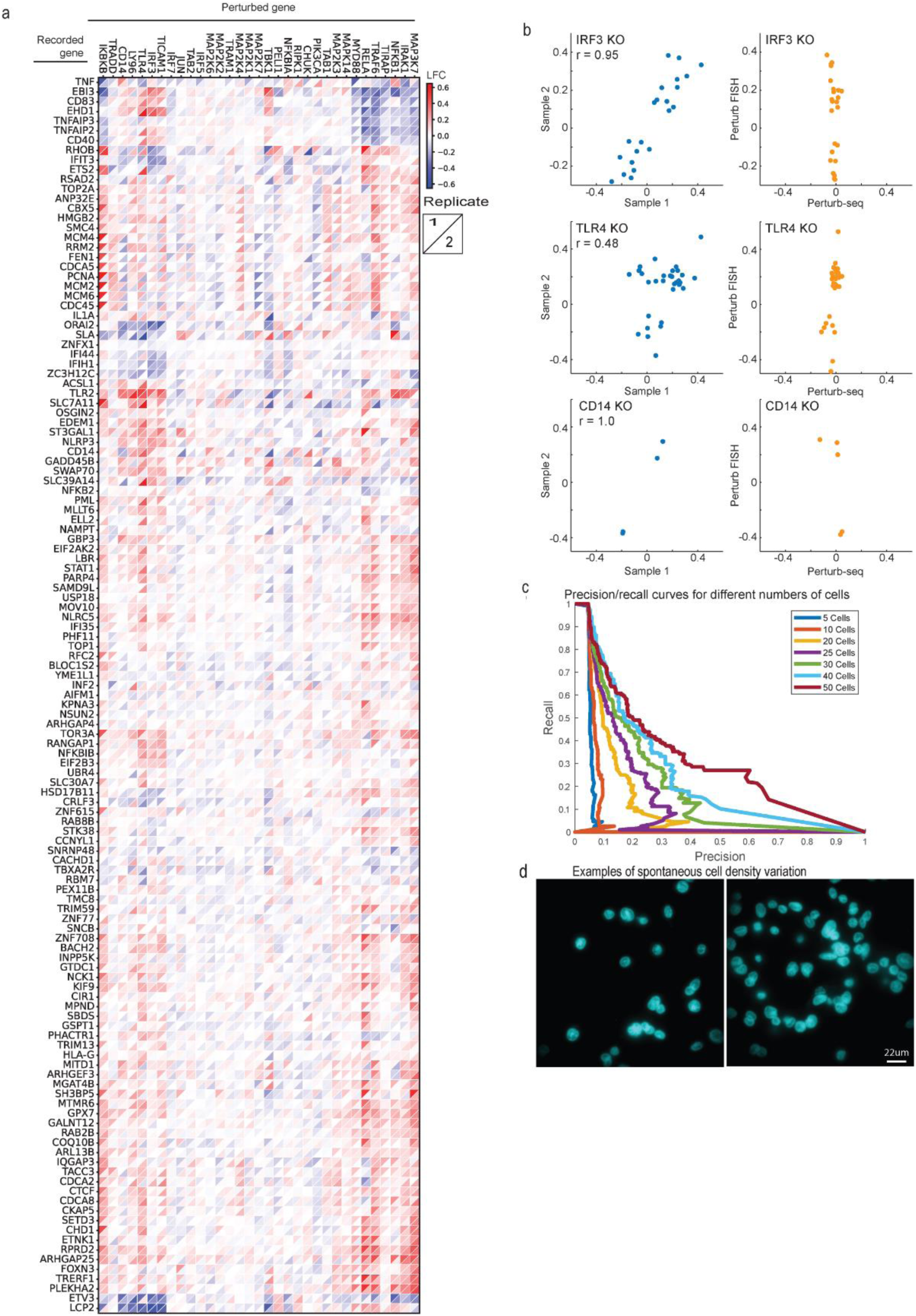
A) Heatmap of LFC effect sizes (colorbar) between Perturb-FISH replicate 1 (upper left triangle of each square) and replicate 2 (lower right triangle). Rows and columns are clustered based on UPGMA clustering of sample1 data. B) Left: scatterplots of significant effect sizes (q<0.1) determined in Perturb-seq (x-axis) and combined Perturb-FISH (y-axis). Right: scatterplots of significant effect sizes (q<0.1) determined in Perturb-FISH replicate 2 (x-axis) and replicate 1 (y-axis). Shown are the effects from IRF3, TLR4 and CD14, which have the lowest correlation between Perturb-FISH and Perturb-seq. Effects represent log-fold changes (LFC; natural log base) in expression relative to control cells. C) Precision recall curves for effects learned in downsampled Perturb-FISH data using increasing numbers of cells. D) Example images (DAPI) of spontaneously occurring areas of varied densities.

**Supplementary figure 2.**
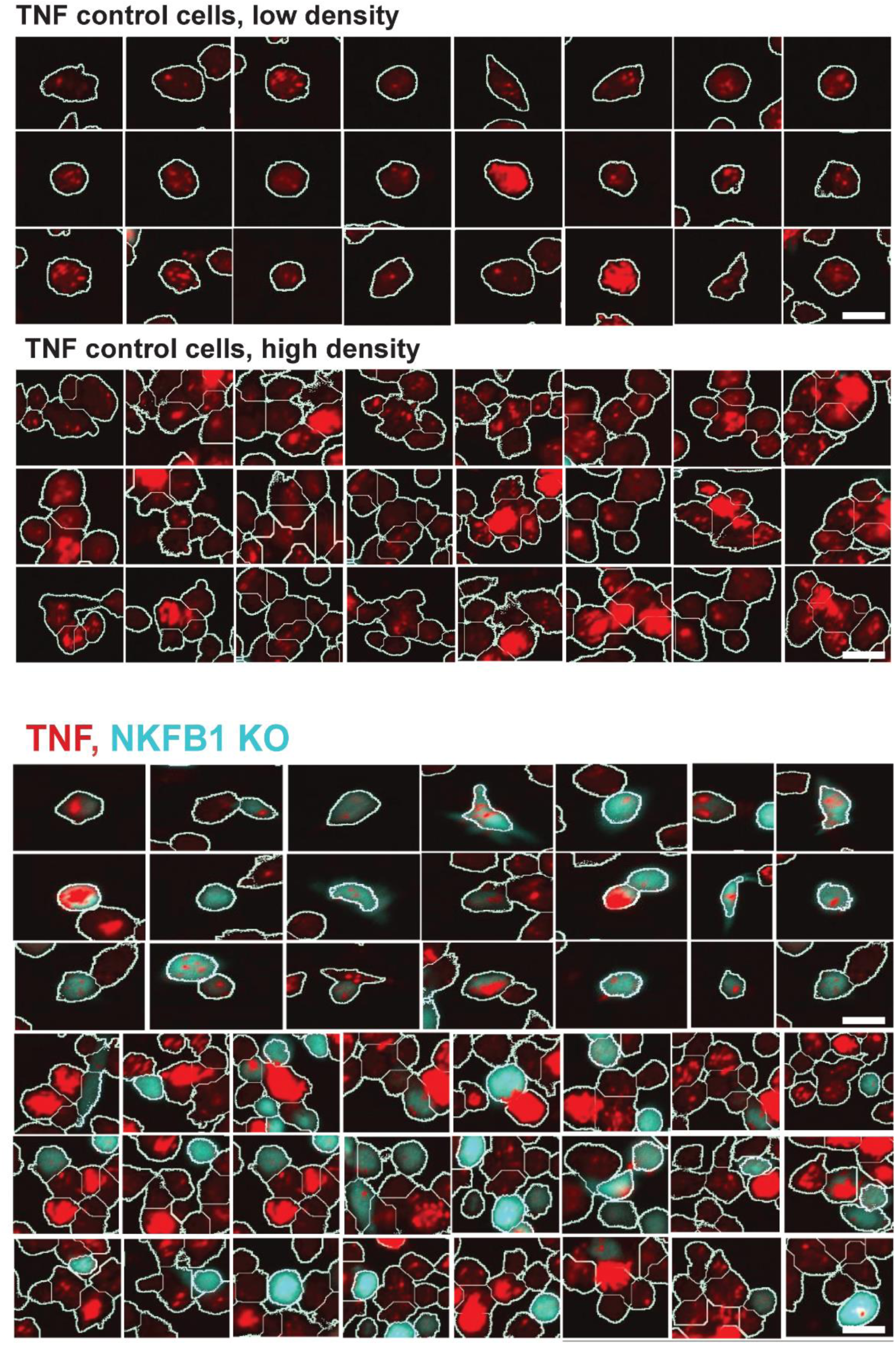
RNAscope images showing TNF mRNAs (in red) in THP1s cells at low and high density. Blue cells received a guide against NFKB1, other cells received a non-targeting guide. Scale bar: 30μm.

**Supplementary figure 3.**
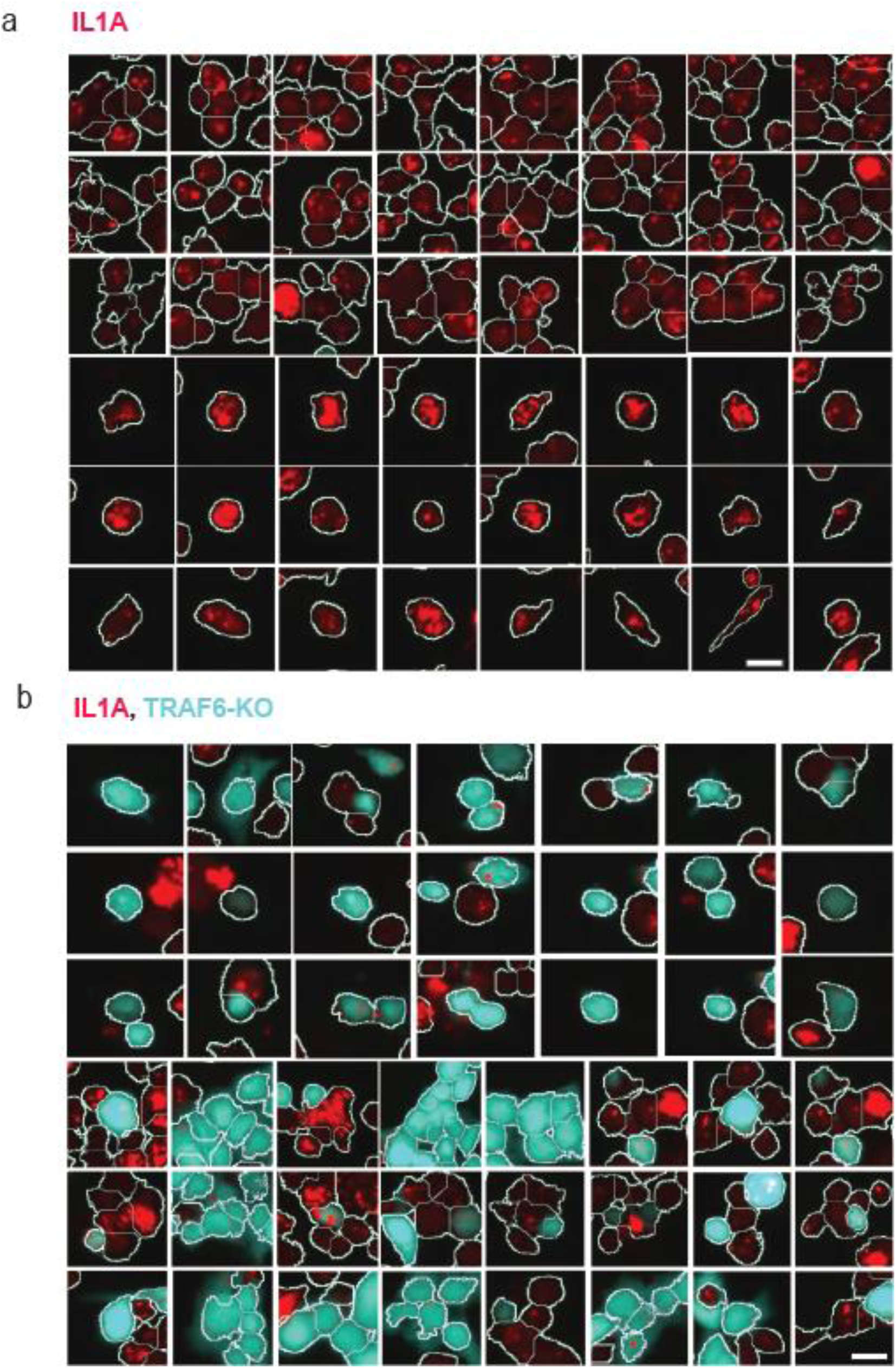
RNAscope images showing IL1a mRNAs (in red) in THP1s at low and high density. Blue cells received a guide against TRAF6, other cells received a non-targeting guide. Scale bar: 30μm.

**Supplementary figure 4.**
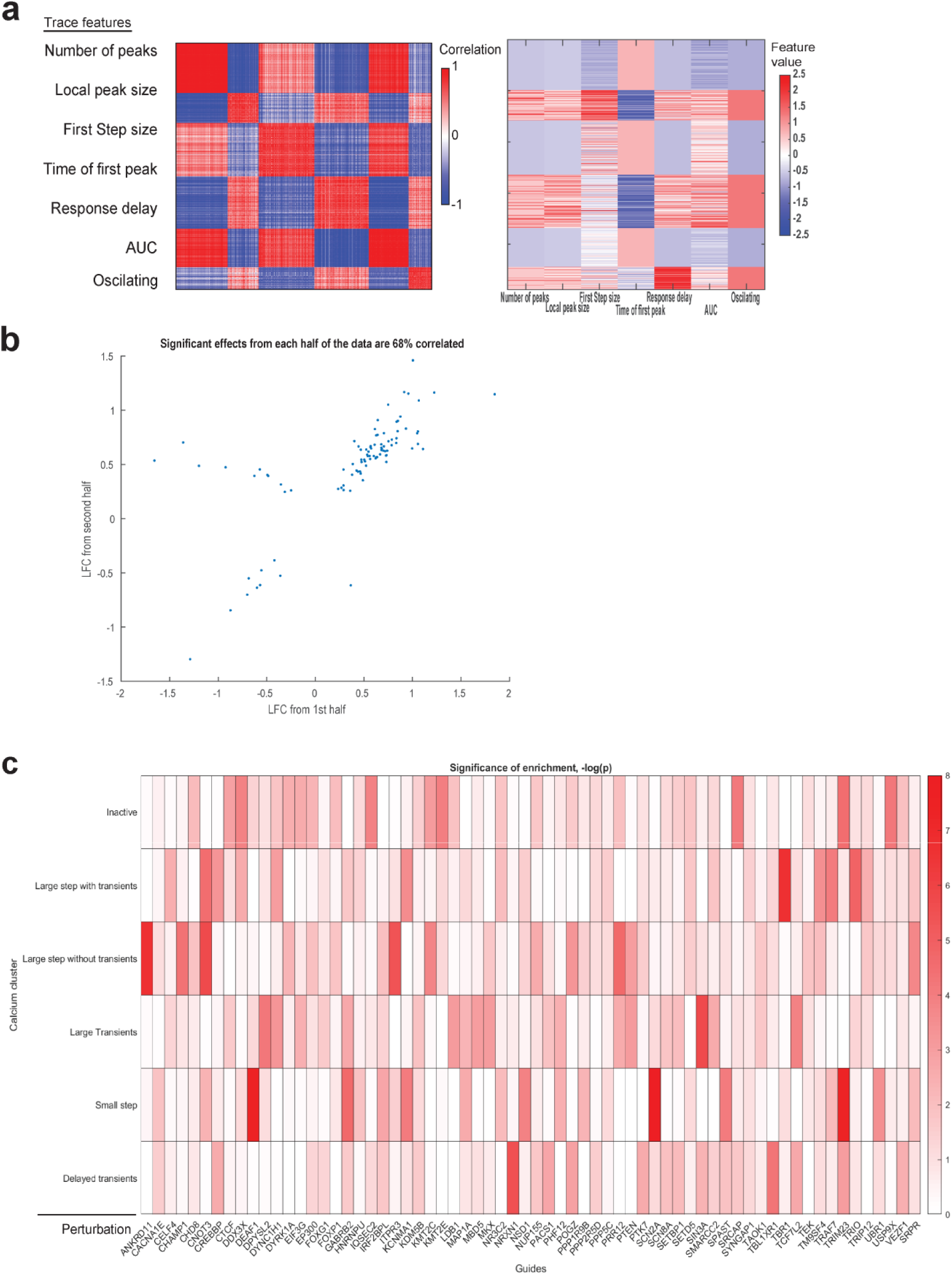
A) Heatmap of cell-to-cell correlation (left) and definition of calcium features (right). Cells are clustered using k-means clustering. B) Scatter plot of significant effects from one half of the data (x-axis) versus the other half (y-axis). C) Heatmap of the significance of the guide (x-axis) enrichment in each cluster. Significance is shown as -log(p-value), using a natural log base. (scale bar).

**Supplementary figure 5.**
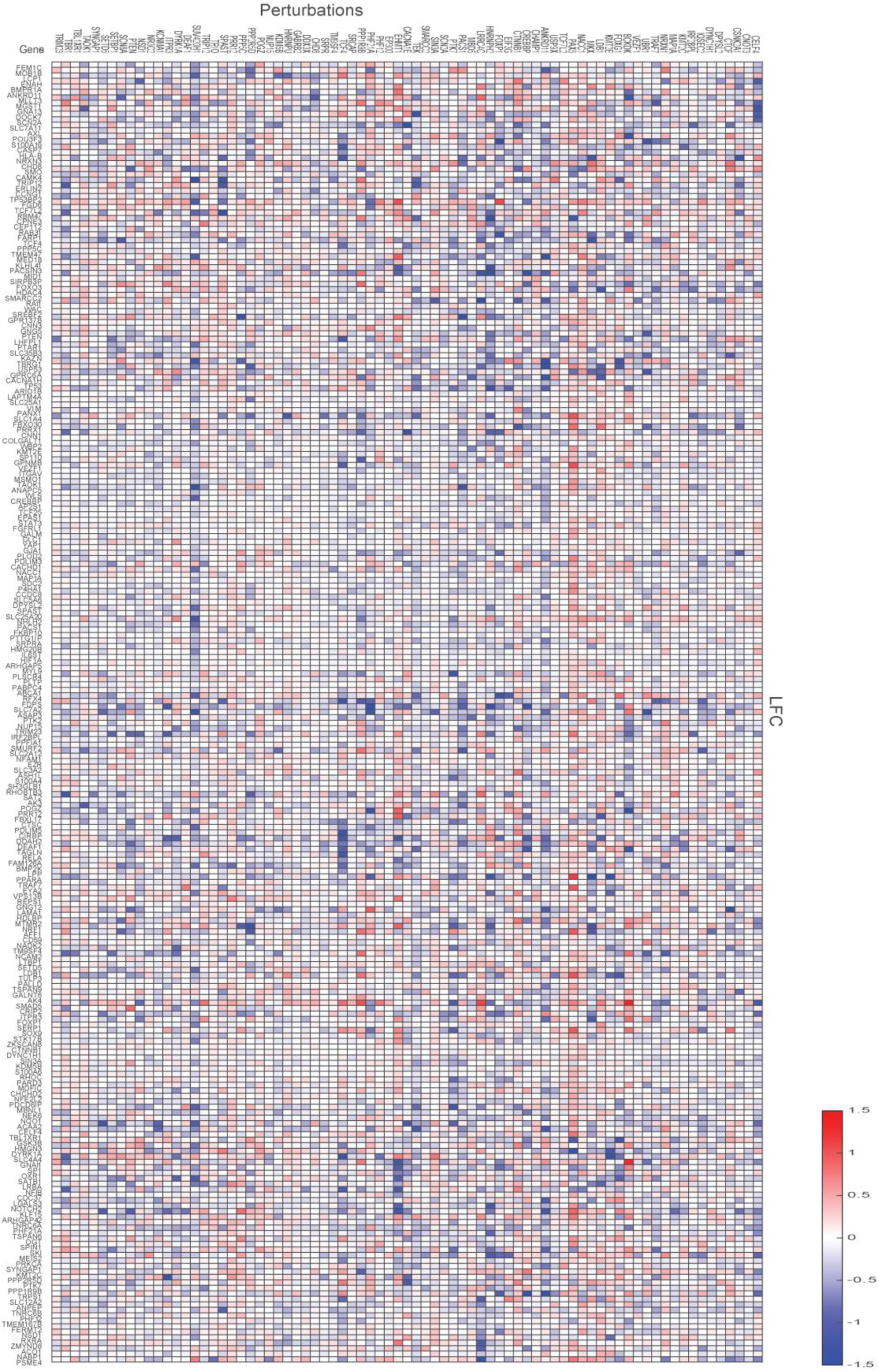
Heatmap showing LFC effects of perturbations (x-axis) on gene expression (y-axis). Effects represent log-fold changes (LFC; natural log base) in expression relative to control cells.

**Supplementary figure 6.**
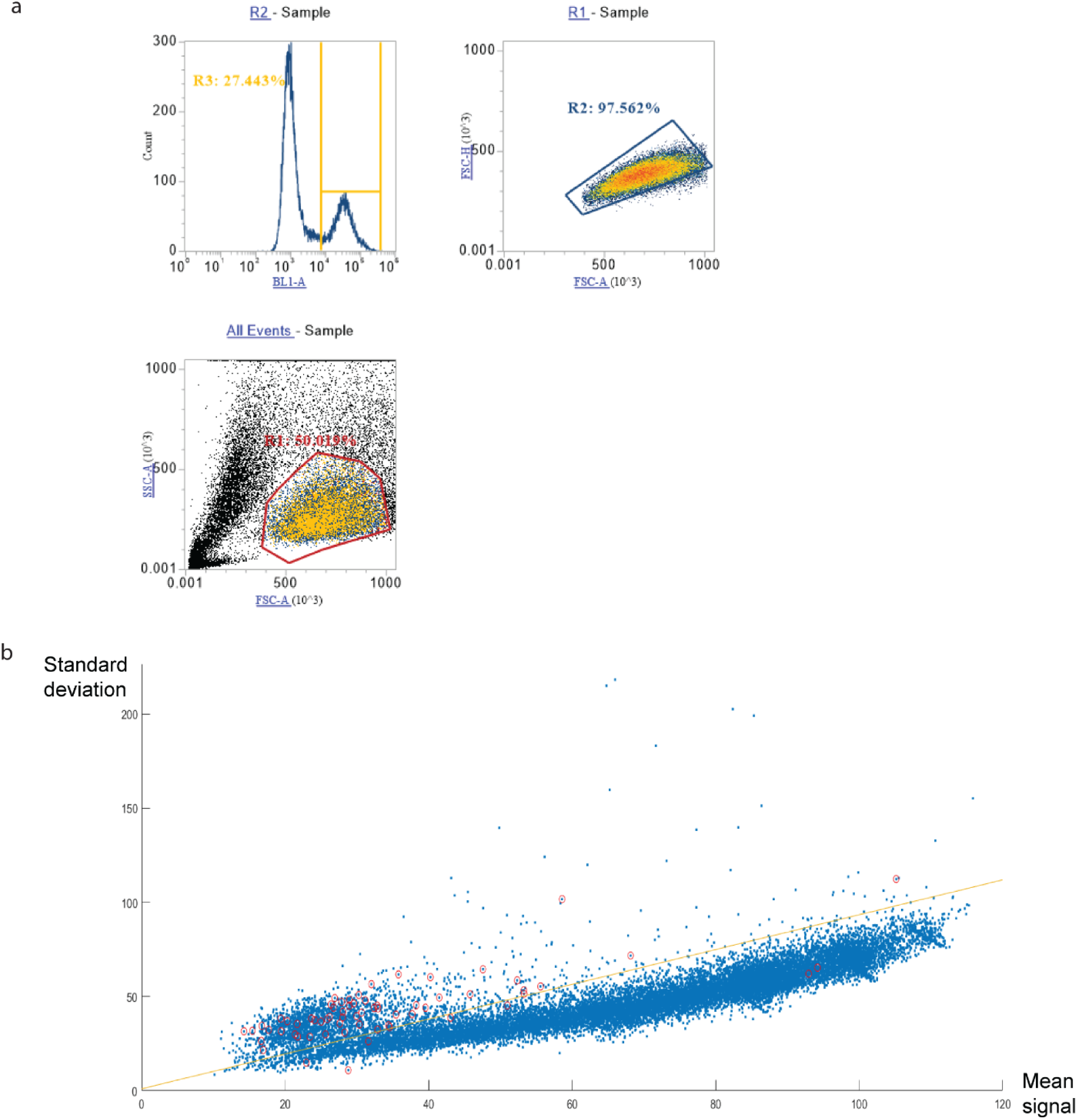
A) Evaluation of Cas9 activity: flow cytometry report of THP1 cells infected with a vector containing both a GFP sequence and a guide against GFP. 73% of the cells are GFP negative (top left). B) Scatter plot of decoded guides showing standard deviation (y-axis) as a function of mean signal across all 15 images (x-axis). Dots circled in red indicate false positives (blank barcodes that do not actually correspond to any guide in the library). The line indicates the applied cut off used during quality filtering of the data. Only decoded guides below the cutoff line were used in downstream analysis.

